# In silico investigation of alternative splicing of microexons in human peripheral tissues

**DOI:** 10.64898/2026.05.13.725022

**Authors:** Shalini Raman, Pallavi Gupta, Ishaan Gupta

**Affiliations:** Indian Institute of Technology Delhi, Department of Biochemical Engineering and Biotechnology, Hauz Khas, New Delhi 110016, India; The University of Queensland Indian Institute of Technology Delhi (UQ-IITD) Research Academy, Hauz Khas, New Delhi 110016, India; The University of Queensland, Australian Institute of Bioengineering and Nanotechnology, St Lucia, QLD 4072, Australia

## Abstract

Microexons are highly conserved fragments of exons ranging from 3 to 51 nucleotides (nts), representing a precise but poorly understood layer of post-transcriptional regulation outside the central nervous system. While their role in neuronal development is well documented, their behavior in peripheral tissues remains largely uncharacterized. In this study, we utilized VAST-TOOLS to perform a comprehensive meta-analysis of alternative microexon splicing across independent transcriptomic datasets spanning hepatic, pulmonary, renal, and colonic tissues. By comparing the diseased and wild-type (WT) profiles, we identified a robust set of differentially spliced microexons (DSMs) unique to each disease. Our findings suggest that microexon dysregulation in the liver, lung, kidney and colon may not be a primary driver of specific diseases, but rather a signature of a broader collapse in cellular splicing homeostasis. We propose that this phenomenon of differential splicing, particularly within critical hub proteins, fundamentally compromises protein interaction networks, thereby priming the cell for the diverse phenotypic failures observed across chronic disease states.

## 1. INTRODUCTION

Alternative splicing (AS) is a highly coordinated, cell type-specific mechanism essential for diverse biological processes, ranging from the maintenance of stem cell pluripotency to the proteomic diversification of the vertebrate tissue (1,2). The fidelity of this process is maintained by the spliceosome, which is a complex ribonucleoprotein machinery that recognizes invariant GU and AG dinucleotides at the 5′ and 3′ splice sites, respectively (3). Consequently, dysregulation of AS is a hallmark of systemic pathology, with aberrant splicing patterns consistently implicated in oncogenesis and neurodegenerative disorders (4,5).

Microexons are defined as the smallest of all exons, typically ranging from 3 to 30 nts, and occasionally up to 51 nt. The splicing of microexons is governed by distinct genomic features. Constitutively spliced microexons possess strong intrinsic signatures that facilitate splicing, including robust splice-site motifs, shorter flanking introns, and a high density of exonic splicing enhancers. In contrast, alternatively spliced microexons are flanked by highly conserved intronic regions harboring specific splicing enhancers. Therefore, their successful inclusion into the final mRNA is driven by RNA motifs bound by helper proteins, such as RBFOX, PTBP1, and SRRM4 (6). Microexons serve as compelling models for AS research in two primary dimensions: first, their minimal size necessitates specialized compensatory enhancing signals for spliceosomal recognition; and second, these brief sequences exert a disproportionately large influence on the functional landscape of the final protein product (7).

The identification of microexons dates back nearly four decades, beginning with the description of 5 nt exons in the *Drosophila Ubx* gene and a 6 nt constitutive microexon in the avian *Troponin T* gene (6). Early investigations into the mammalian brain further highlighted the prevalence of these events, such as the developmentally regulated 30 nt microexon in rat *Ncam* and the strikingly small 3 nt microexon in the same gene identified in mouse brain (8). Foundational studies establish microexons as precisely timed regulatory elements with dramatic inclusion shifts during central nervous system (CNS) differentiation. For instance, as mouse embryonic stem cells become glutamatergic neurons, nearly 70% of regulated microexons show >50% inclusion changes. This includes the 12-nt microexon in *Enah* gene (9), which shifts from embryonic exclusion to high mature-neuron inclusion to modulate actin assembly. Although comprising only ∼1% of all AS events (4), microexons account for roughly one-third of conserved, isoform-generating neural AS events. Furthermore, the deep conservation of intronic sequences flanking these in-frame microexons, spanning 450 million years of vertebrate evolution, underscores their vital functional roles and the evolutionary pressure to maintain their regulatory architecture (8). In coding regions, the impact of microexon inclusion is largely dictated by the reading frame. Approximately 80-90% of microexons are in-frame, allowing them to subtly modify protein-protein interactions, enzymatic activity, or sub-cellular localisation without disrupting the overall protein fold (10). Conversely, out-of-frame “poison” microexons introduce premature stop codons, triggering nonsense-mediated decay (NMD) to fine-tune steady-state gene expression, as seen in the inflammation-linked transcription factor *NFKB1* (6). Despite this extensive characterisation within the CNS, the behaviour of microexons in peripheral metabolic and respiratory tissues remains comparatively obscure, limited to that of constitutive microexons across these tissues as well as their low basal inclusion levels being altered in tumors (10). We focus our analyses on the lung, kidney, colon, and liver, which are among the highly alternatively spliced organs (11). We mapped alternative splicing events by comparing diseased transcriptomes to their respective WT controls. Specifically, our pulmonary analysis was focussed at four distinct datasets: Small Cell Lung Cancer (SCLC), Idiopathic Pulmonary Fibrosis (IPF), Chronic Obstructive Pulmonary Disease (COPD), and COVID-19. The hepatic investigation covered pathologies like Chronic Hepatitis (CH), Liver Fibrosis (LC), Dysplastic Nodules (DN), alongside early and advanced Hepatocellular Carcinoma (HCC). Furthermore, the renal and colonic systems were evaluated through two targeted conditions each: Diabetic Nephropathy (DNE) and Renal Cell Carcinoma (RCC) in the kidney, alongside Ulcerative Colitis (UC) and Colorectal Cancer (CC) in the colon. Through this multi-organ approach, we delineate tissue-specific splicing signatures and elucidate the broader role of transcriptomic dysregulation in chronic diseases.

## 2. RESULTS

### 2.1. Characterising microexon diversity across wild-type human peripheral tissues

We begin by performing a cross-cohort study on distinct lung pathology datasets to set a baseline trend of the AS of microexons in peripheral tissues. We analyzed the WT controls from cohorts dedicated to IPF, SCLC, COPD, and COVID-19, comprising 19, 7, 88, and 10 samples respectively. Using the VAST-TOOLS framework, we detected microexon splicing across three size-based categories: 1 -15 nts, 16 - 27 nts, and 28 - 51 nts. Their percent spliced-in (PSI) values were calculated to indicate the proportion of reads that support their inclusion across all reads supporting the splice site pair (12). While VAST-TOOLS outputs all AS events, including alternative exons, alternative splice-site usage, and retained introns, we focus on 16976 events that can be classified as microexons according to the mentioned categories (**Supplementary Fig. 1**).

Our analysis revealed that despite the varied origins of these samples, the pulmonary baseline spliced-in microexons remained remarkably consistent [**Fig**. **1**]. Specifically, between 27% and 56% of all microexons were detected across all four independent datasets, while 63 - 83% were shared in at least three. This high degree of overlap demonstrates a stable core of microexon inclusion in healthy lung tissue. The distribution of these events, contrasted against the total number of annotated events in the VAST-DB, is detailed in **table 1**, highlighting the subset of total microexons that are active in the pulmonary tissue.

**Fig 1:**
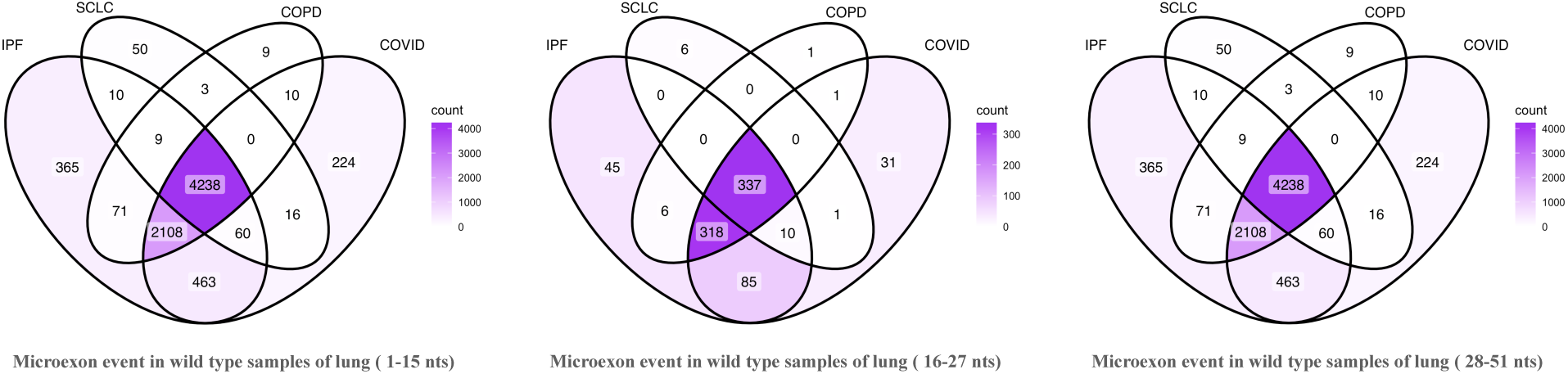
Major overlap between microexon events that occur exclusively in the lung tissues;

**Table 1:** Total no. Of microexons in each category.

| Dataset | No. of microexon event occurrences in the inclusion file, according to the VASTDB database | Microexon events specific to control dataset |  |  | Cutoffs taken for filtering |  |
| --- | --- | --- | --- | --- | --- | --- |
|  |  | upto 15 nts | 16 - 27 nts | 28 - 51 nts | min_samples | psi_threshold |
| SCLC wildtype | Upto 15 nts : 2356<br>16 - 27 nts : 1707<br>28 - 51 nts : 12913 | 85 | 354 | 4386 | 2/7 | > $\pm 10\%$ |
| COPD wildtype |  | 171 | 663 | 6448 | 27/88 |  |
| IPF wildtype |  | 250 | 801 | 7324 | 6/19 |  |
| COVID wildtype |  | 231 | 783 | 7119 | 3/10 |  |

Having established a robust and consistent microexon baseline across multiple cohorts, we expand our analysis to include WT samples from liver, kidney, and colon. We benchmark these against the brain, which is the primary standard for microexon studies (11), and aim to delineate a core non-neuronal splicing program which is highly understudied in the past research concerning microexons.

Our multi-organ comparison reveals that microexon regulation in these peripheral sites is distinct and not merely a subset of the brain’s program [**Fig. 2a**]. While the brain maintains the highest number of tissue-specific microexons, the lung appears next with a unique subset, followed by a smaller number of colon-, kidney-, and liver-specific microexons. There also exists a large number of microexons shared across two or more tissues. This matches the previously established canonical trend of the proportion of genes with skipped exons in various tissues (11). We also find that each of the tissues had increasing proportions of microexons across the three size-based bins, with the 28-51 nt category emerging as the most dominant category [**Fig. 2b**]. This observation demands checking if and how the organ-specific microexon inclusion is disrupted across diseases.

**Fig 2:**
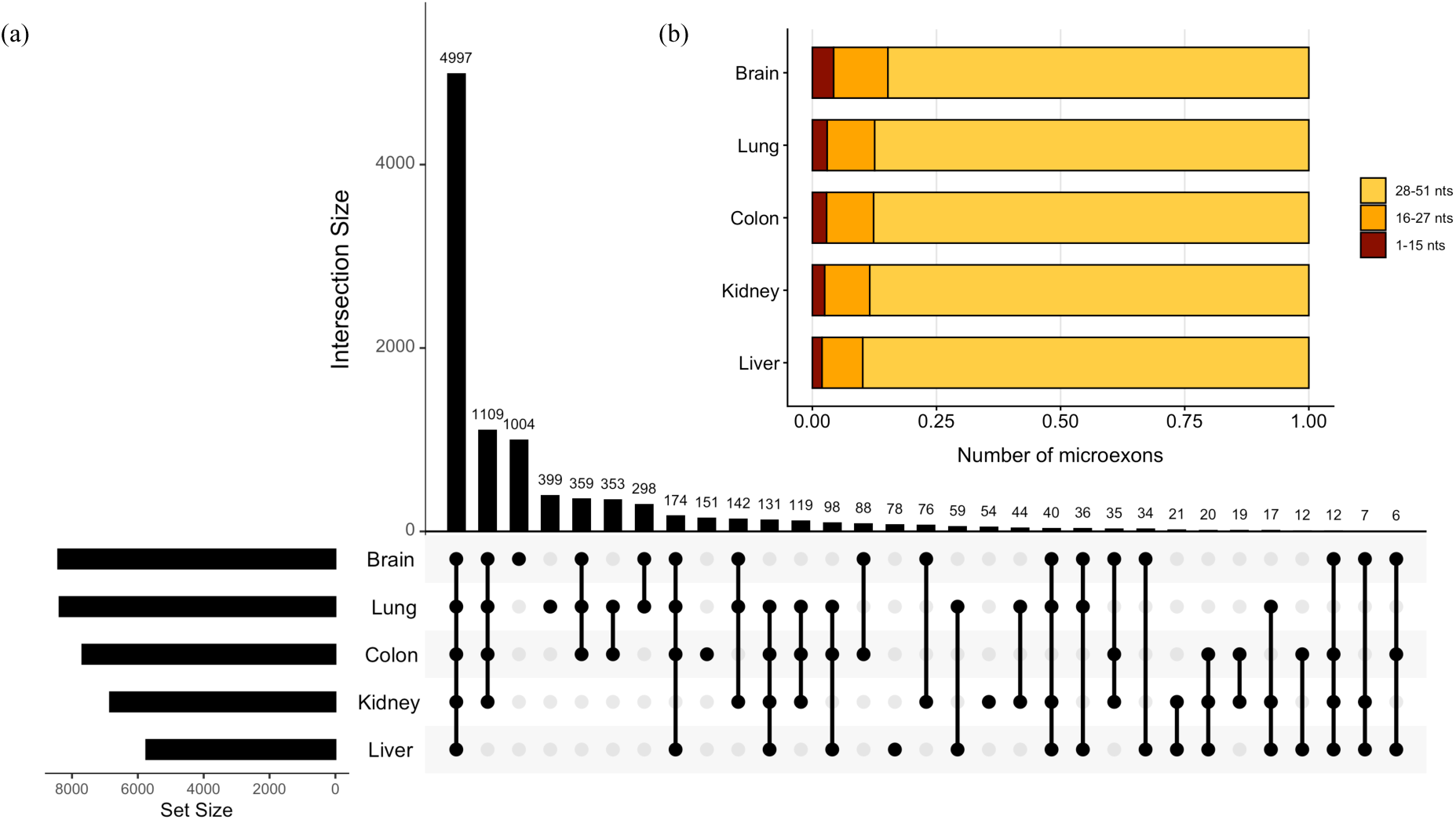
Intersecting microexon events occurring in wild type samples of brain, lung, colon, kidney and liver tissues. (a) Plot showing the intersection of the microexons upto a length of 51 nts in all the respective tissue samples; (b) Plot showing the distribution of the microexons ranging in three different categories based on length in wild type conditions

### 2.2. Quantifying aberrant microexon inclusion patterns across multiple peripheral diseases

We begin by checking disruptions in microexon splicing in lung pathologies, including SCLC (Control: 7; Patient: 79), IPF (Control: 19; Patient: 20), COPD (Control: 88; Patient: 93), and COVID-19 (Control: 10; Patient: 26). We observe aberrant splicing across each of these diseases, encompassing microexons from all size ranges [**Fig. 3a** and **Table 2**]. Approximately 81-91% of all mis-spliced microexons, irrespective of the size-based tier, were specific to the individual etiologies, with COVID-19 demonstrating the highest number, followed by IPF. This length-independent degree of dysregulation proves that this non-canonical splicing of short yet significant transcriptomic sequences observed in viral, fibrotic, and cancerous conditions is driven by independent, cause-specific RNA profiles rather than a generalised or shared pulmonary stress response. This shift in the regulatory logic across different lung conditions suggests that while the baseline state of a particular tissue can be uniform, the transition to disease can be inherently stochastic and divergent. Unlike the previous reports from the post-mortem autism brain (13), where increased microexon skipping was observed, we find both increase and decrease in their inclusion in each of the lung diseases [**Fig. 3b-e**].

**Fig. 3:**
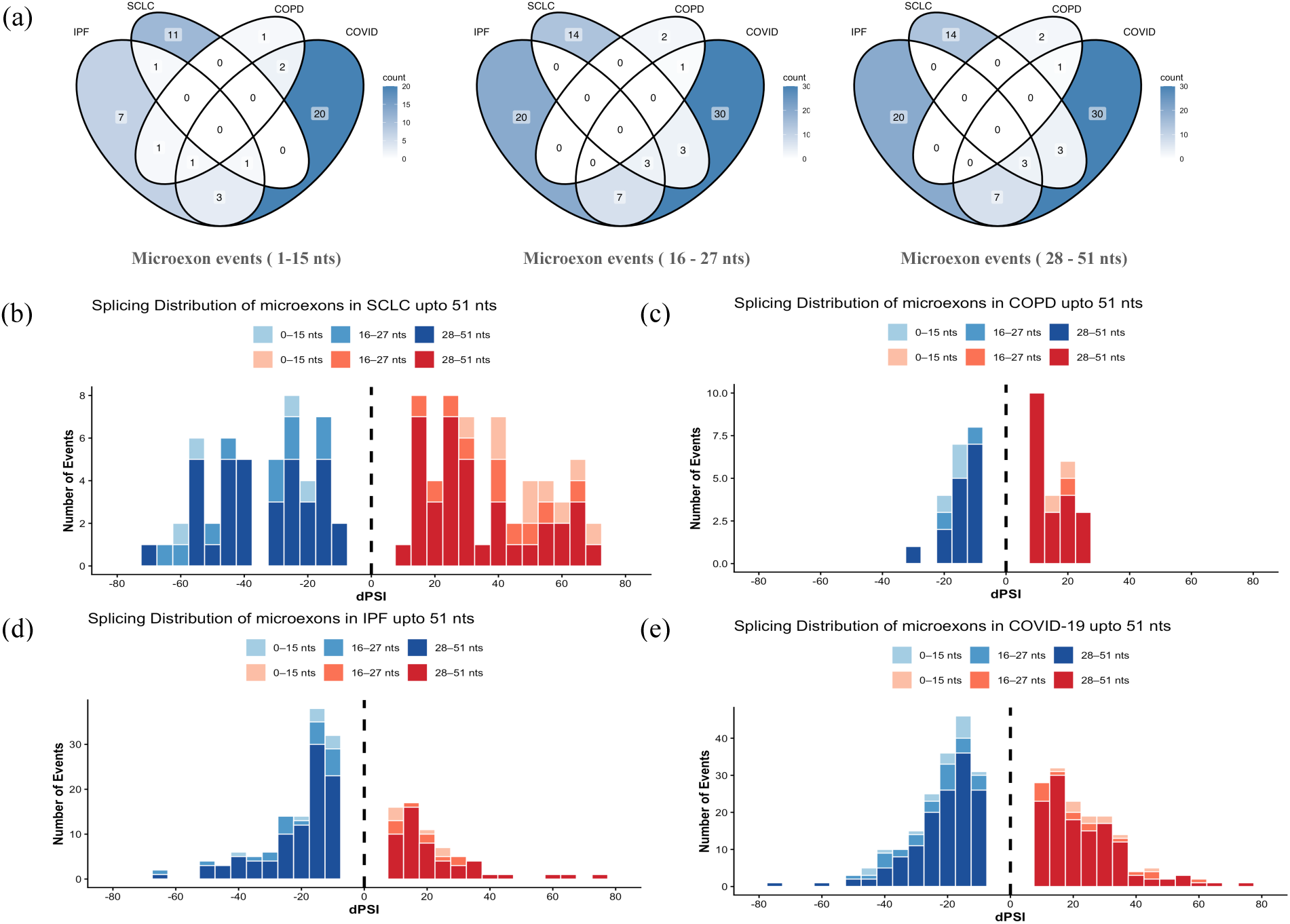
(a) No. of DSMs occurring in lung diseases; (b-e) Histograms depicting the distribution of dPSI values of microexons ranging upto 51 nts in length, where blue bars indicate decreased inclusion and red indicate increased inclusion, for the diseases IPF, SCLC, COPD and COVID respectively.

**Table 2:** No. of microexon events classified on the basis of their inclusion levels in the respective disease conditions and categories based on length in lung

| Disease | MIC events (1-15 nts) |  | MIC events (16-27 nts) |  | MIC events (28-51 nts) |  |
| --- | --- | --- | --- | --- | --- | --- |
| PSI values | Increasing | Decreasing | Increasing | Decreasing | Increasing | Decreasing |
| SCLC | 9 | 4 | 11 | 13 | 47 | 50 |
| IPF | 6 | 8 | 9 | 21 | 50 | 95 |
| COPD | 2 | 3 | 2 | 2 | 23 | 15 |
| COVID | 11 | 15 | 15 | 27 | 118 | 133 |

Continuing the analyses, we checked two pathologies each for the human colon and kidney, namely UC (Control:15;Patient:30) and CC (Control:18;Patient:18) for the former and DNE (Control:9;Patient:27) and RCC (Control:7;Patient:27) for the latter. As was seen for lung diseases, those tested here shared almost negligible DSMs [**Fig. 4a** and **Fig. 5a**, respectively]. Within this rigidly isolated framework in the colon, UC consistently dominates the differential event count, driving approximately 60-68% of the unique splicing shifts across all size tiers, while CC independently accounts for roughly 30-35%. For the kidney, across all microexon length categories, a remarkable ∼95% of identified splicing alterations remain strictly exclusive to either DNE or RCC. Each of the tested pathologies of the colon and the kidney, whose WT microexon abundances are second to that of the lung, are correlated and often coexist. For example, studies have reported increased risk of CC in patients with UC (14); however, ∼95% of all identified microexon splicing alterations remain strictly exclusive to a single disease state in all the three length categories. Similarly, although patients with diabetes are at a significantly increased risk of developing DNE and RCC (15) they share a negligible number of mis-spliced microexons. Such unique transcriptomic signatures could potentially serve as novel biomarkers to otherwise overlapping pathologies. Moreover, this disruption too occurred in both directions with several microexons showing increased inclusion and others with increased skipping [**Fig. 4b-c** and **Table 3** and **Fig. 5b-c** and **Table 4**, respectively].

**Fig. 4:**
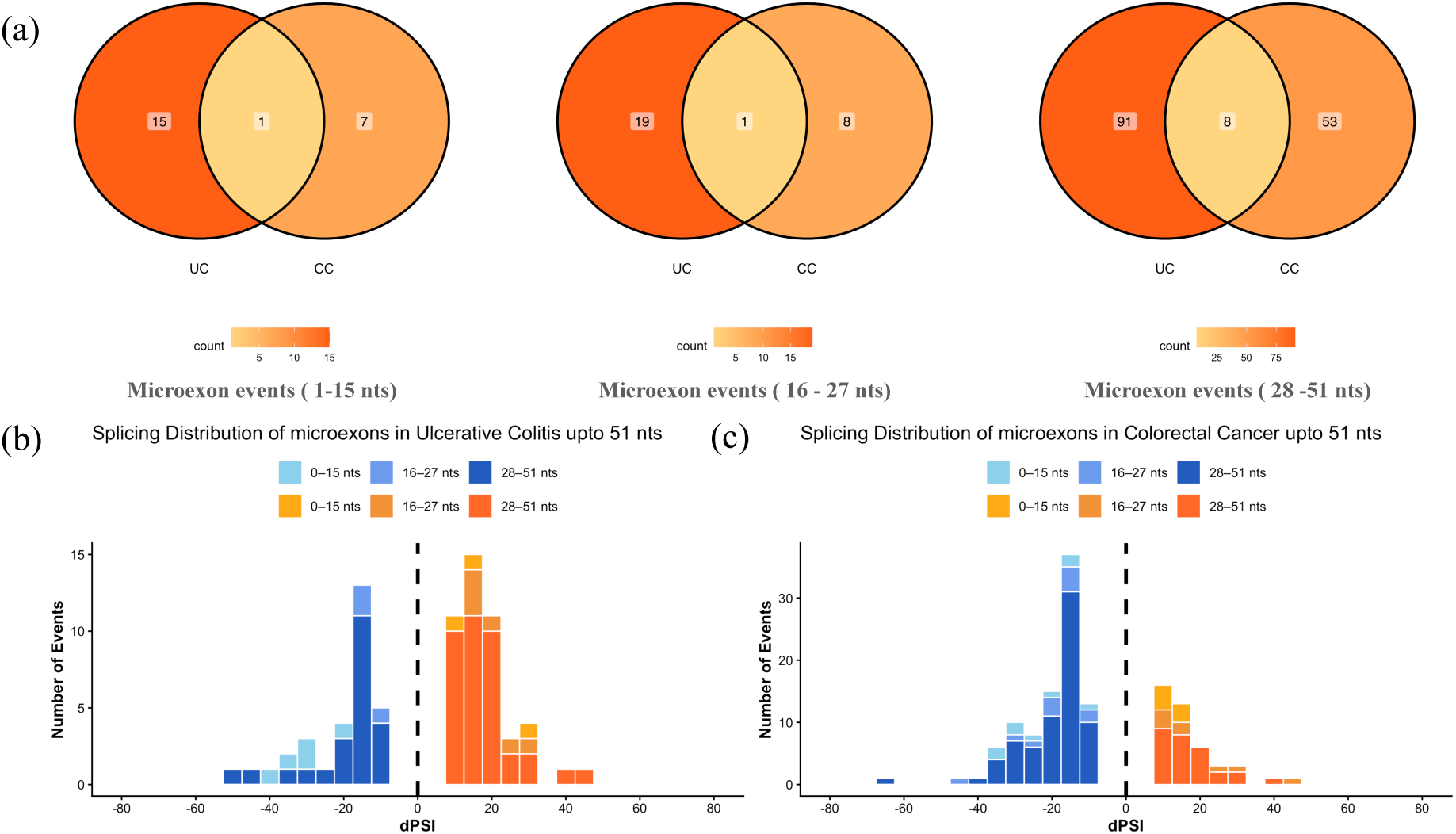
(a) No. of DSMs occurring in colon diseases; (b&c) Histograms depicting the distribution of dPSI values of microexons ranging upto 51 nts in length, where blue bars indicate decreased inclusion and orange indicate increased inclusion for the diseases UC and CC.

**Fig. 5:**
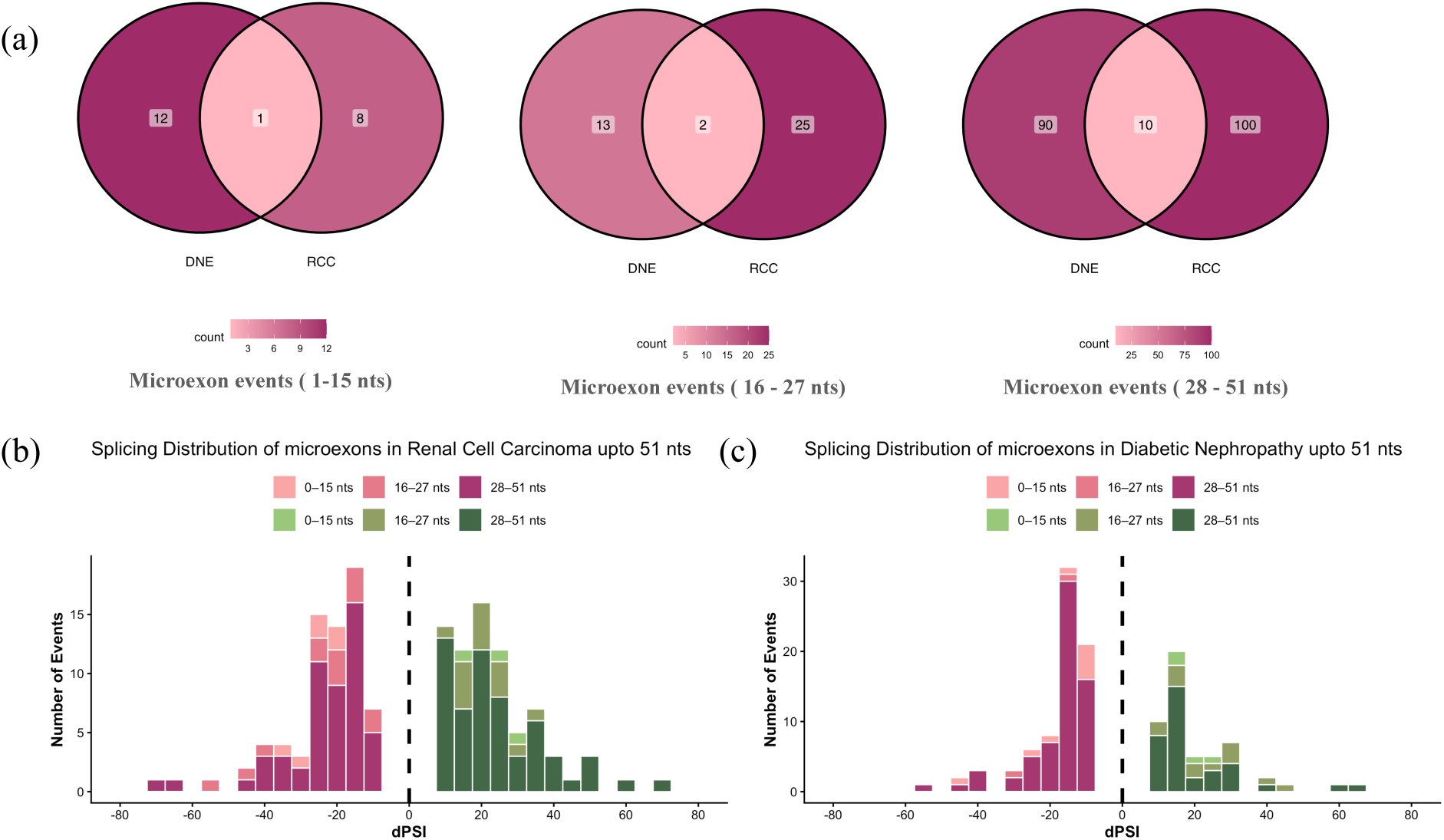
(a) No. of DSMs occuring in kidney diseases; (b&c) Histograms depicting the distribution of dPSI values of microexons ranging upto 51 nts in length, where dark pink bars indicate decreased inclusion and green indicate increased inclusion for the diseases RCC and DNE.

**Table 3:** No. of microexon events classified on the basis of their inclusion levels in the respective disease conditions and categories based on length in colon

| Disease | MIC events (1-15 nts) |  | MIC events (16-27 nts) |  | MIC events (28-51 nts) |  |
| --- | --- | --- | --- | --- | --- | --- |
| PSI values | Increasing | Decreasing | Increasing | Decreasing | Increasing | Decreasing |
| CC | 7 | 9 | 8 | 12 | 28 | 71 |
| UC | 3 | 5 | 6 | 3 | 37 | 24 |

**Table 4:** No. of microexon events classified on the basis of their inclusion levels in the respective disease conditions and categories based on length in kidney

| Disease | MIC events (1-15 nts) |  | MIC events (16-27 nts) |  | MIC events (28-51 nts) |  |
| --- | --- | --- | --- | --- | --- | --- |
|  | Increasing | Decreasing | Increasing | Decreasing | Increasing | Decreasing |
| RCC | 3 | 6 | 14 | 13 | 58 | 52 |
| DNE | 4 | 9 | 13 | 2 | 35 | 65 |

The final evaluation was carried out on liver diseases: NAFLD (Control:7;Patient:10), HCC, LC, DN & CH (diseases other than NAFLD in the liver cohort were from the same study with 13 control samples and 46, 8, 5, and 12 for HCC, LC, DN & CH respectively) [**Fig. 6**][**Table 5**]. The liver is amongst the least alternatively spliced tissues in humans (11), which is reflected in the relatively low number of DSMs found across the tested liver pathologies [**Fig. 6a and Table 5**]. It is noteworthy that both early and advanced HCC have both common and unique disruptions to microexon inclusion with the latter showing a larger number, as expected owing to a more advanced disease stage. Again, all the tested diseases have microexons with both positive and negative ΔPSI values [**Fig. 6b-g**].

**Fig. 6:**
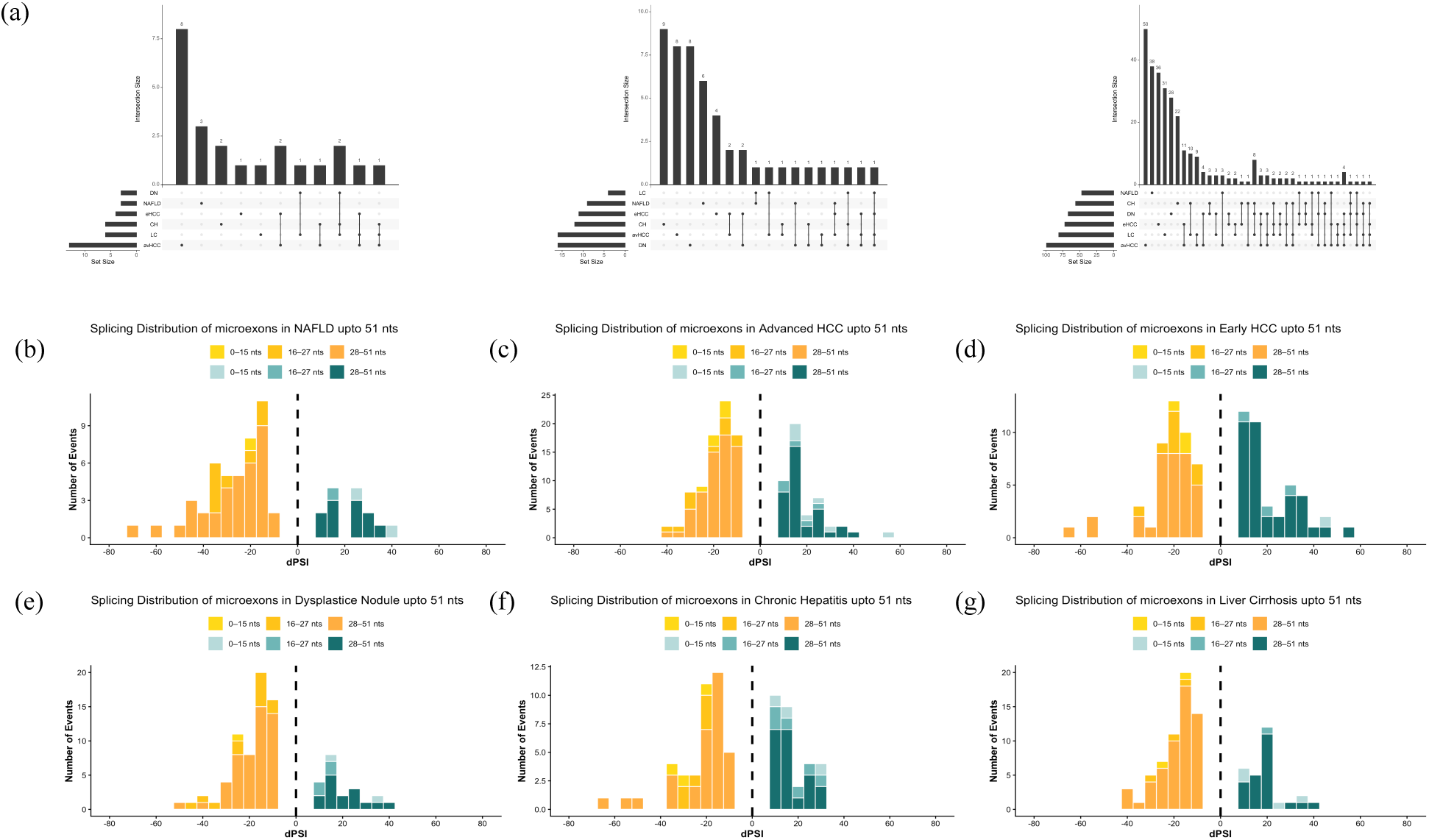
(a) No. of DSMs occuring in liver diseases; (b-g) Histograms depicting the distribution of dPSI values of microexons ranging upto 51 nts in length, where brown bars indicate decreased inclusion and yellow indicate increased inclusion for the diseases NAFLD, HCC, DN, CH and LC.

**Table 5:** No. of microexon events classified on the basis of their inclusion levels in the respective disease conditions and categories based on length in liver

| Disease | MIC events (1-15 nts) |  | MIC events (16-27 nts) |  | MIC events (28-51 nts) |  |
| --- | --- | --- | --- | --- | --- | --- |
| PSI values | Increasing | Decreasing | Increasing | Decreasing | Increasing | Decreasing |
| avHCC | 7 | 6 | 5 | 11 | 35 | 64 |
| eHCC | 1 | 3 | 3 | 8 | 37 | 35 |
| Chronic Hepatitis | 3 | 3 | 6 | 6 | 20 | 36 |
| Dysplastic Nodule | 1 | 2 | 4 | 12 | 16 | 51 |
| Liver Cirrhosis | 4 | 2 | 1 | 3 | 24 | 57 |
| NAFLD | 2 | 1 | 1 | 8 | 11 | 36 |

### 2.3. Microexon misregulation impacts genes linked to disease pathology

Although we find a small number of highly DSMs in the tested diseases, there exists a potential for them to play a significant role by altering the host gene’s expression as well as cause alterations in the resultant protein function or protein interaction networks. Below we discuss some of the genes that show microexon |ΔPSI| ≥ 20% as well as have a previously reported role in the respective disease pathology (**Supplementary Table 1**).

#### 2.3.1 Genes in Pulmonary Diseases

Several DSMs in SCLC occur in genes linked to the disease and tissue dysfunction. For instance, DSM inclusion was increased in *AP1S2* [**Fig. 7a**] encodes for a Golgi protein which recruits clathrin, recognizes sorting signals, and is a miR-204-5p target regulating lung cancer behavior (16). Increased inclusion was also found in a microexon in *PACSIN2* [**Fig. 7b**], a kinase substrate identified as an extracellular vesicular proteomic signature in lung cancer plasma (17), and in PKCα, whose interaction with *DLG1* correlates with increased invasiveness (18). Conversely, DSM inclusion was decreased in *TMEM87A* [**Fig. 7c**] and *CLIP1*, genes forming critical fusions (*TMEM87A-RASGRF1* and *CLIP1-LTK*) that act as potential biomarkers and therapeutic targets in lung cancer (19,20).

**Fig. 7:**
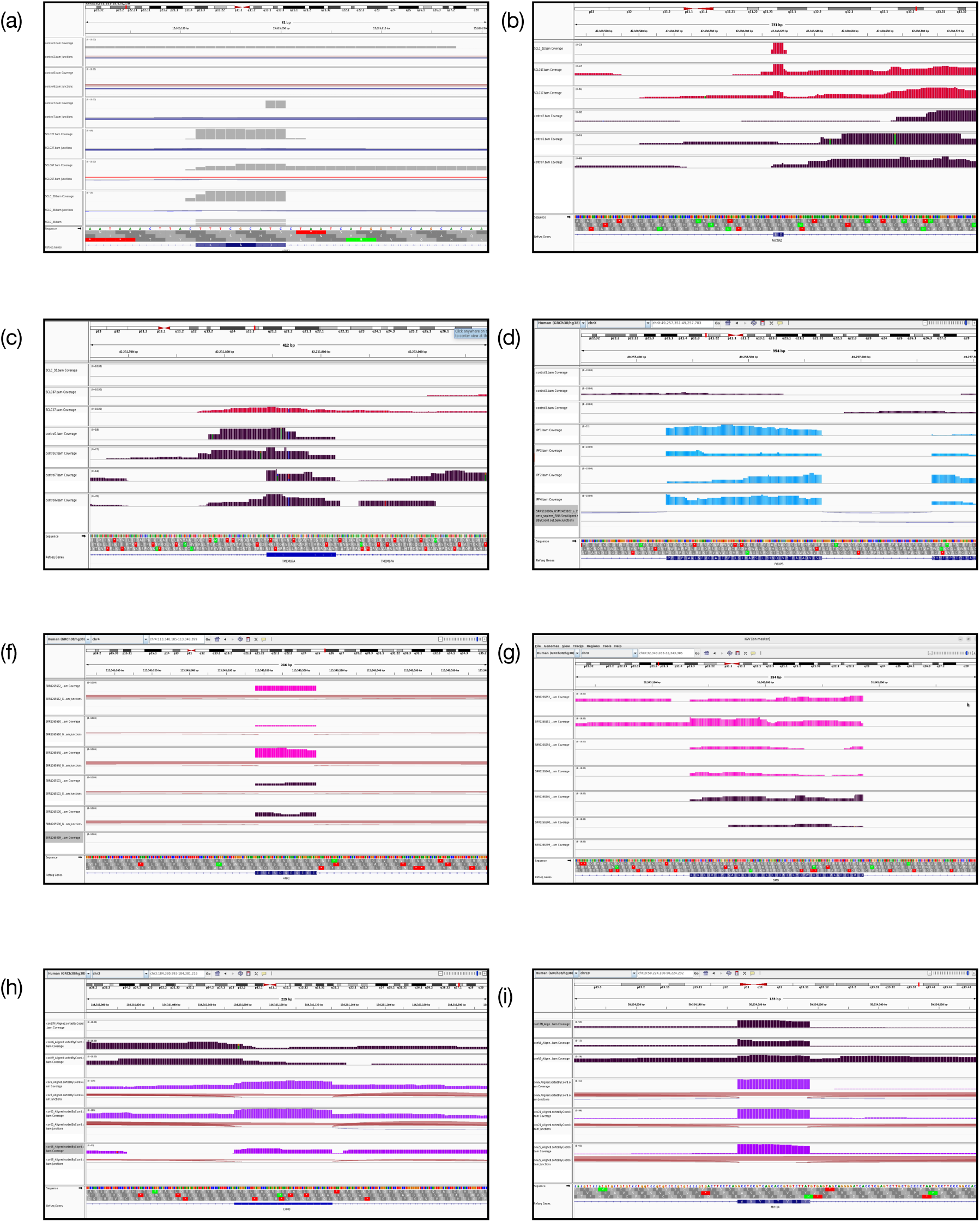
IGV snapshots of Differentially Spliced Microexons (DSMs) across multiple disease datasets. The panels illustrate varying levels of inclusion for specific genes: (a–c) SCLC dataset: (a) *AP1S2* (increased inclusion), (b) *PACSIN2* (increased inclusion), and (c) *TMEM87A* (decreased inclusion). Red coverage tracks/ markings indicate SCLC patient samples, while dark tracks indicate controls. (d) IPF dataset: *FOXP3* showing increased inclusion. Blue coverage tracks indicate IPF patient samples, and dark tracks indicate controls. (e–f) COPD dataset: (e) *ANK2* (increased inclusion) and (f) *DMD* (increased inclusion). Pink coverage tracks indicate COPD patient samples, and dark tracks indicate controls. (g–h) COVID-19 dataset: (g) *CHRD* (increased inclusion) and (h) *MYH14* (increased inclusion). Purple coverage tracks indicate COVID-19 patient samples, and dark tracks indicate controls. All dark-colored coverage tracks across panels represent respective control samples for each dataset.

In IPF, decreased DSM inclusion occurs in *CPNE7* (Copine7), encoding calcium-dependent membrane-binding proteins that predict disease severity (21), and in *HNF4G* (Hepatocyte Nuclear Factor 4 Gamma). While not directly linked to IPF, *HNF4G* downregulation via miR-320b suppresses lung cancer growth and angiogenesis (22), highlighting an increased lung cancer risk in fibrosis patients (23). Conversely, increased DSM inclusion was observed in *ADAM28* (24), *TMEM71* (Transmembrane Protein 71) (25), and *FOXP3* (Forkhead box P3) (26) [**Fig. 7d**], genes closely associated with lung cancer when overexpressed.

The COPD microexon profile reveals links to Lung adenocarcinoma (LUAD) and IPF, aligning with known COPD-LUAD coexistence (27) and IPF commonalities (28). Increased DSM inclusion occurs in *ANK2* [**Fig. 7e**] , a potential biomarker for immune checkpoint inhibitor (ICI) efficacy in LUAD (29), and in *DMD* [**Fig. 7f**] , matching its differential expression in IPF (30). Conversely, decreased microexon inclusion was found in *NCAM1*, whose expression is altered by a smoking-related genetic variant driving COPD (31). Interestingly, while *EGFLAM* showed decreased inclusion in our analysis, other studies report its elevated expression in COPD patients (32).

In COVID-19, characterized by complex viral entry mechanisms, immune suppression, and structural respiratory failure, increased DSM inclusion occurs in *NCAM1*, *CHRD* (Chordin) [**Fig. 7g**], and *MYH14* (Myosin Heavy Chain 14) [**Fig. 7h**], genes closely associated with the disease (33–35). Conversely, decreased inclusion was observed in *CTC1* and *FSD1L*. Though not directly linked to COVID-19, they play significant roles in NSCLC and lung squamous cell cancer (LSCC), respectively (36,37)

#### 2.3.2 Genes in Colon Diseases

Analysis of colon diseases like CC and UC revealed critically misregulated genes. In CC, increased DSM inclusion occurred in *COLQ*, whose genetic variants link to colonic structural conditions like diverticulitis (38), and in *DMD*, a biomarker predicting early-onset colorectal cancer development and prognosis via the miR-31-5p-DMD axis (39). Decreased inclusion was found in a microexon in the lncRNA *TONSL-AS1*, heavily implicated in gastric cancer progression (40) and relevant here due to gastric-colon cancer synchrony (41), as well as in *cTAGE5*, a tumor-associated antigen frequently detected in colon carcinoma (42).

In UC, increased DSM inclusion was observed in microexons in *ALG11* and *ZNF226*, both known contributors to colonic pathologies like cancer (43,44). Conversely, decreased inclusion was found for a DSM in *ZIK1*, a transcriptional repressor linked to intestinal metaplasia and chronic tissue injury (45), and in *CCNE1*, whose notable elevation in early tumor stages significantly correlates with overall colon cancer survival (46).

#### 2.3.3 Genes in Kidney Diseases

The differential analysis of renal pathologies revealed some critically significant genes that have been reported for playing a role in the respective diseases. In RCC, increased DSM inclusion was observed in *XKR8*, which encodes a phospholipid scramblase facilitating efferocytosis.This altered splicing may disrupt normal activity and reinforce an immunosuppressive tumor microenvironment (47). Increased inclusion also occurred in *GGA3*, an endosomal adaptor whose altered form enhances receptor recycling to fuel the aggressive, invasive phenotype of renal carcinomas (48). Conversely, decreased inclusion was found in *BAZ2B*, whose mRNA expression is significantly lower in clear cell and chromophobe RCCs (49), and in *ZBTB14*, a transcription factor whose motifs enrich hypomethylated enhancers during kidney cancer development (50).

In DNE, microexon splicing may be predictive of poorer prognosis; for instance, increased inclusion was observed in *COG1*, linked to unfavorable survival in KIRC (51), and *CPNE7*, a strong predictor of end-stage renal disease (ESRD) (52). Decreased inclusion was identified in a microexon in *SCGB2B2*; while its upregulation is heavily linked to proteinuria and podocyte dysfunction marking DNE progression (53,54), our analysis suggests its decreased DSM inclusion yields a structurally deficient isoform. Similarly, decreased inclusion (negative ΔPSI) in *ADA* emphasizes its functional contribution to metabolic and structural disruptions in Diabetic Kidney Disease (55).

#### 2.3.4 Genes in Liver Diseases

Our study encompasses some of the most common diseases that occur in the liver tissues and we have uncovered some very important genes with DSMs playing an important role in the malignancies associated with the liver in various capacities. Because HCC accounts for 85-90% of all liver cancers (56), identifying specific genes and splicing events behind its pathogenesis is crucial for developing targeted diagnostics. In early HCC, increased DSM inclusion was found in *NDUFAF2*, an elevated independent prognostic biomarker (57), and *USP21* [**Fig. 8d**], which promotes proliferation alongside GTF3C2 upregulation (58). Conversely, decreased inclusion was observed in *TRERF1* (Transcriptional Regulating Factor 1), an HBV-targeted gene differentially expressed in cholangiocarcinoma (59), and *MAP3K9* (Mitogen-Activated Protein Kinase Kinase Kinase 9), which is differentially expressed in HCC (60). In advanced HCC, the splicing program shifts toward aggressive malignancy and treatment failure. Increased inclusion was seen in *PRUNE2*, linked to HCC mutations (61), and in *FAM13A* [**Fig. 8a**], which promotes Cisplatin-resistant cell proliferation (62). In contrast, decreased inclusion occurred in *AP3S1* [**Fig. 8c**], an overexpressed prognostic biomarker across multiple cancers including HCC (63), and in *USO1* [**Fig. 8b**], whose isoforms dysregulate the endoplasmic reticulum-Golgi network to promote liver cancer progression (64).

**Fig. 8:**
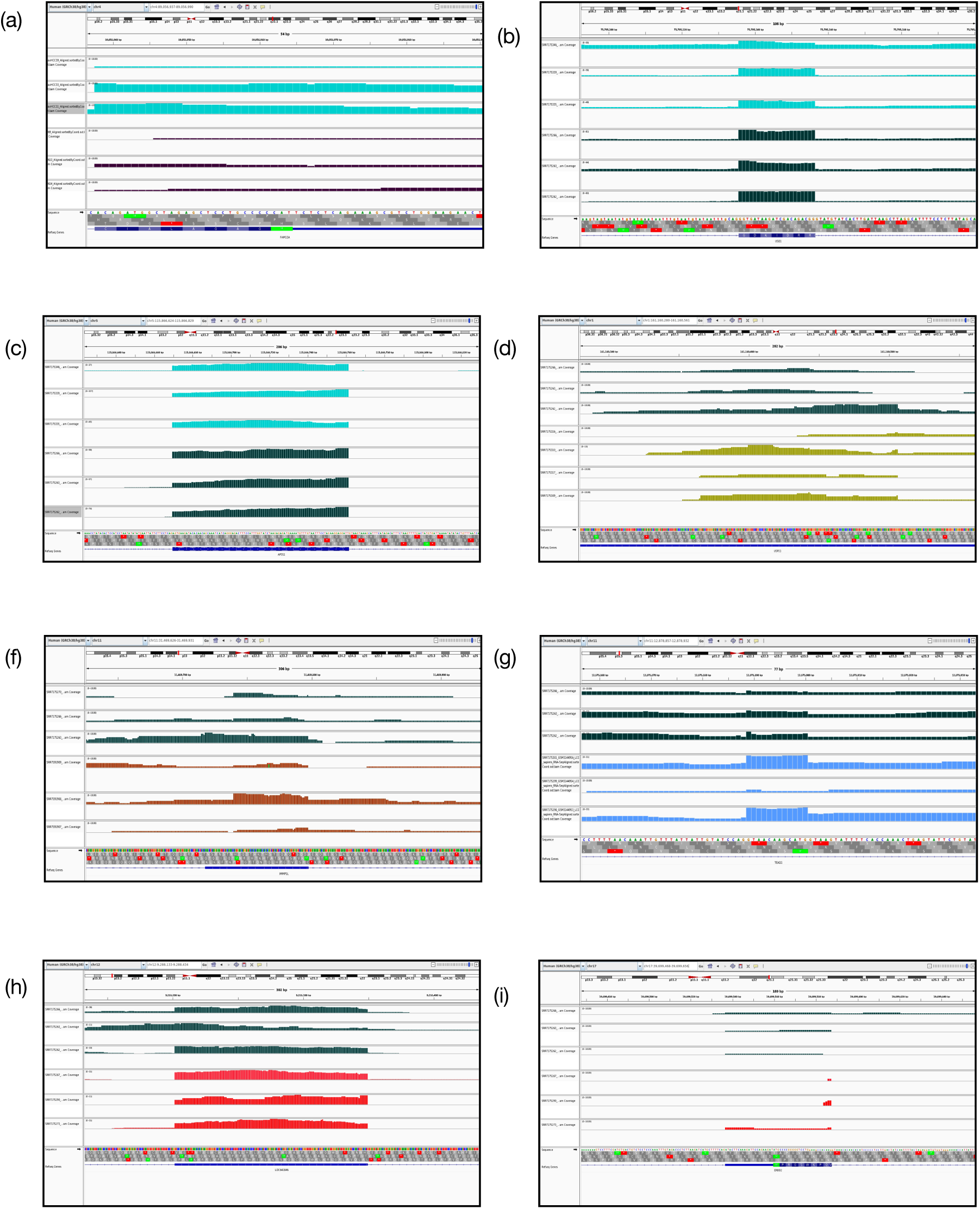
IGV snapshots of Differentially Spliced Microexons (DSMs) in HCC, DN, LC, and CH datasets. Panels represent varied inclusion patterns across tissues: (a–d) HCC dataset: (a) *FAM13A* (increased inclusion), (b) *USO1* (decreased inclusion), (c) *AP3S1* (decreased inclusion), and (d) *USP21* (increased inclusion). Dark coverage tracks in (a–c) represent advanced HCC controls, while dark tracks in (d) represent early HCC controls. (e) DN dataset: *IMMP1L* (increased inclusion, ΔPSI = 81.03). (f) LC dataset: *TEAD1* (increased inclusion, ΔPSI = 24.61450549.. (g–h) CH dataset: (g) *DDX11* (increased inclusion) and (h) *ERBB2* (decreased inclusion). Red coverage tracks indicate CH patient samples, while dark tracks indicate respective controls. Unless otherwise specified, dark-colored tracks across all panels represent control samples for each condition.

In DN, a precancerous HCC stage, altered splicing highlights early malignant rewiring (65). Genes with increased DSM inclusion help establish foundational tumor networks. For instance, protein-protein interaction networks reveal that *IMMP1L* [**Fig. 8e**] co-expresses with *DNAJC24* [**Fig. 9d**], a potential HCC therapeutic target, strongly linking this interaction to the DN-to-HCC transition (66). While not previously defined as direct DN drivers, the identified genes are heavily implicated in HCC pathology. Increased inclusion also occurs in *NACC1*, featured in hepatic carcinoma-specific fusions (67), and *UPF3B*, which accelerates liver cancer proliferation via CDK12-mediated autophagy (68). Conversely, decreased inclusion was found in *ZBTB14*, whose role in promoting NAFLD-associated fibrosis (69) suggests its dysregulation facilitates the shift to nodular dysplasia. Similarly, decreased inclusion in *MAP3K6* correlates with its downregulated mRNA expression in cryptogenic HCC (70).

**Fig. 9:**
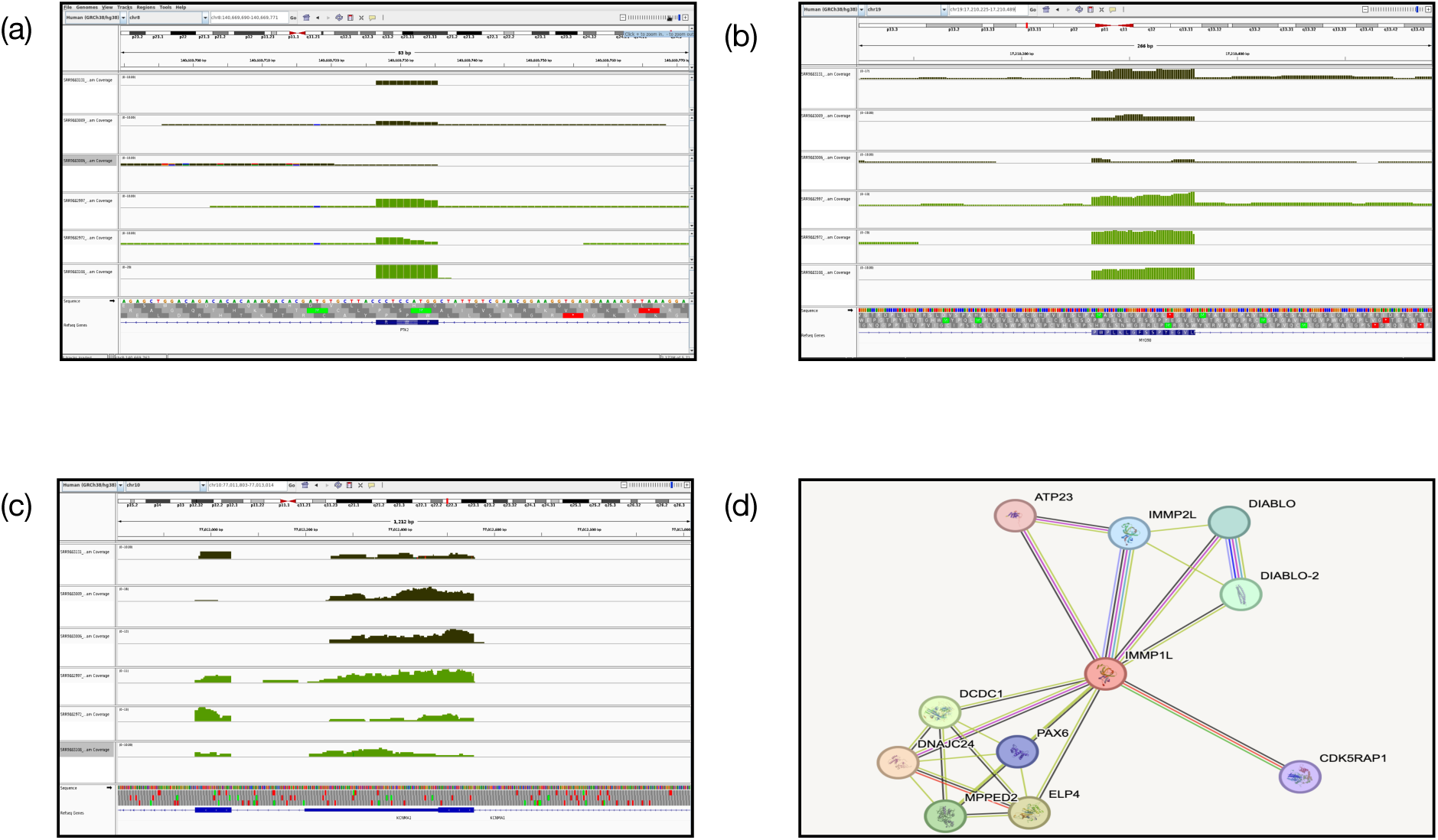
IGV snapshots of DSMs in the NAFLD dataset and protein-protein interaction network. (a–c) NAFLD dataset: IGV snapshots showing different levels of inclusion for (a) *PTK2* (increased inclusion), (b) *MYO9B* (increased inclusion), and (c) *KCNMA1* (decreased inclusion). Green coverage tracks indicate NAFLD patient samples, while dark-colored tracks indicate respective controls. (d) Protein interaction network (STRING database) illustrating the predicted interactions between *IMMP1L* and *DNAJC24*.

Liver cirrhosis (LC) indicates persistent injury overlapping with HCC, NAFLD, and Hepatitis. During LC, increased DSM inclusion often drives systemic stress responses and metabolic remodeling. For example, increased inclusion occurs in *BAZ2B*, a broad cancer prognostic marker (including LIHC) whose splicing alteration may precede malignancy (49). Similarly, increased inclusion was found in *TEAD1* [**Fig. 8f**], whose expression is remodeled by estrogen receptors to alleviate hepatic steatosis, a cirrhosis precursor (71). Conversely, decreased inclusion was observed in *GUCY1A2*, which shares promoter methylation status with HBV-related HCC (72), and in *ALPK3*, linked to increased HCC cell viability and stemness (73).

In chronic hepatitis (CH), DSMs reflect an interplay between viral infection and early oncogenic activation, correlating with progression to severe liver pathologies. Increased DSM inclusion occurs in *KSR1* and *DDX11* [**Fig. 8g**], both linked to core signaling dysregulation in HCC (74,75). Because CH is a major cause of HCC (76,77), exploring these early transcriptomic alterations is crucial to solidify the molecular link between chronic inflammation and malignancy. Conversely, decreased inclusion was found in viral adaptation genes like *ERBB2* [**Fig. 8h**], associated with Hepatitis C prevalence (78), and *PRC1*, which silences protective genes via chromatin compaction to facilitate metabolic rewiring and liver malignancy (79).

In NAFLD, increased DSM inclusion affects complex axes modulating disease severity. This includes *PTK2* [**Fig. 9a**], a marker of active metabolic reorganization though its circular RNA suppresses NAFLD via the miR-200c/SIK2/PI3K/Akt axis (80), and *MYO9B* [**Fig. 9b**], which modulates metabolic transition to non-alcoholic steatohepatitis (NASH) via miR-193b-5p (81). In contrast, our analysis indicates that microexon skipping in genes such as *DDX11* and *KCNMA1* [**Fig. 9c**] which represent an early loss of regulatory control, potentially contributing to disrupted metabolic homeostasis and oncogenic suppression in studies (82) and (83).

## 3. DISCUSSION

The purpose of this analysis was to explore the relevance of non-neuronal microexons, which is currently a class of genomic features that is significantly understudied in ongoing transcriptomic research. While literature confirms that the brain remains the most active environment for alternative splicing of exons (including microexons), we find, in our evaluated peripheral tissues, namely lung, colon, kidney, and liver, that they show varying basal microexon inclusion [**Fig. 2b**], as suggested previously for exons (11). However, it was unknown if microexons are mis-spliced in these, otherwise highly, alternatively spliced tissues. After setting a precedent in the baseline trends of microexon splicing in WT tissues, we observed a contrasting difference between the WT stability and the pathological divergence of microexons. Across all the tested WT and diseased tissues, the aberrant splicing does not follow a singular, predictable trajectory, but rather reflects a localized breakdown of regulatory precision unique to each etiology.

Genes harbouring dramatic positive or negative ΔPSI values were back-traced to further document their association with the progression of their respective disease pathologies. By explicitly detailing the interplay of these genes in the respective conditions, this study highlights that microexons can serve as pivotal markers of the disease state and suggests that the non-neuronal tissues hold untapped potential for understanding the molecular architecture of several diseases. Therefore, as suggested previously, these minimal yet significant cases of differential splicing require more attention and exploration to explain the molecular phenotype of these etiologies better.

## 4. METHODS

### 4.1 Data Acquisition

Publicly available bulk RNA-Seq data for all the disease cohorts was downloaded from NCBI repositories like GEO (Gene Expression Omnibus). Data for the pulmonary diseases: SCLC, IPF, COPD and COVID 19 was acquired from four different studies: PRJNA257389; PRJNA358081, PRJNA24581 & GSE183533. Data for renal and colonic diseases: DNE, RCC, CC & UC were acquired from different studies: PRJNA595590; PRJNA704974; PRJNA218851 and PRJNA1019284. Finally, data for the hepatic cohort was chosen from two different studies: PRJNA471801 and PRJNA558102.

### 4.2 Data Pre-processing

Md5sum was checked for all the files downloaded. For each analysis, the FASTQ reads were pre-processed, trimmed and de-duplicated to avoid bias across the samples using fastqc (v 0.12.1) (https://github.com/s-andrews/fastqc), and additionally, MultiQC (v 1.31) (84) and FASTp (v 1.0.1) (85) to ensure consideration of good quality data for further analysis.

### 4.3 Bioinformatics analysis

VAST-TOOLS (Vertebrate Alternative Splicing and Transcription Tools) (https://github.com/ vastgroup/vast-tools) was employed to identify all the annotated and hypothetical splice sites as well as their resulting alternative exons (AltEx), retained introns (IR), and alternative 5’ (donor) and 3’ (acceptor) splice site usage (Alt5/ Alt3) (https://github.com/vastgroup/vast-tools). The baseline trend of inclusion of microexons in WT samples was calculated by identifying microexon events in expressed genes by applying a PSI cutoff of > ±10 in at least 30% of the samples considered. For the differential splicing analysis, ΔPSI values were calculated to quantify the shifts in microexon inclusion levels across the two different conditions (control vs patient). To ensure statistical robustness and biological relevance, we employed the Wilcoxon test for the calculation due to the high number of samples per dataset. A stringent filtering cutoff was applied, as in the case of WT analysis, where a microexon was only considered for analysis if at least 30% of the samples in a given comparison group contained non-NA values. Furthermore, we defined the significance threshold for differential inclusion using a ΔPSI cutoff of ΔPSI > 10% and p.val < 0.05, categorizing events as increased or decreased based on whether the inclusion level was more than 10% or less than -10%, respectively.

To validate the DSM findings from the VAST-TOOLS output, the read alignment of those genes containing the DSMs was observed in IGV (86) to visualize the difference in the levels of the expression of the respective genes. Bam files were generated for various samples using the STAR (Spliced Transcripts Alignment to a Reference) tool (87) to serve as input for the IGV software. All the plots were made using R programming by the virtue of several packages like ggplot2, Euler and UpSetR.

