## Supplementary Fig. 1 for "In silico investigation of alternative splicing of microexons in human peripheral tissues"

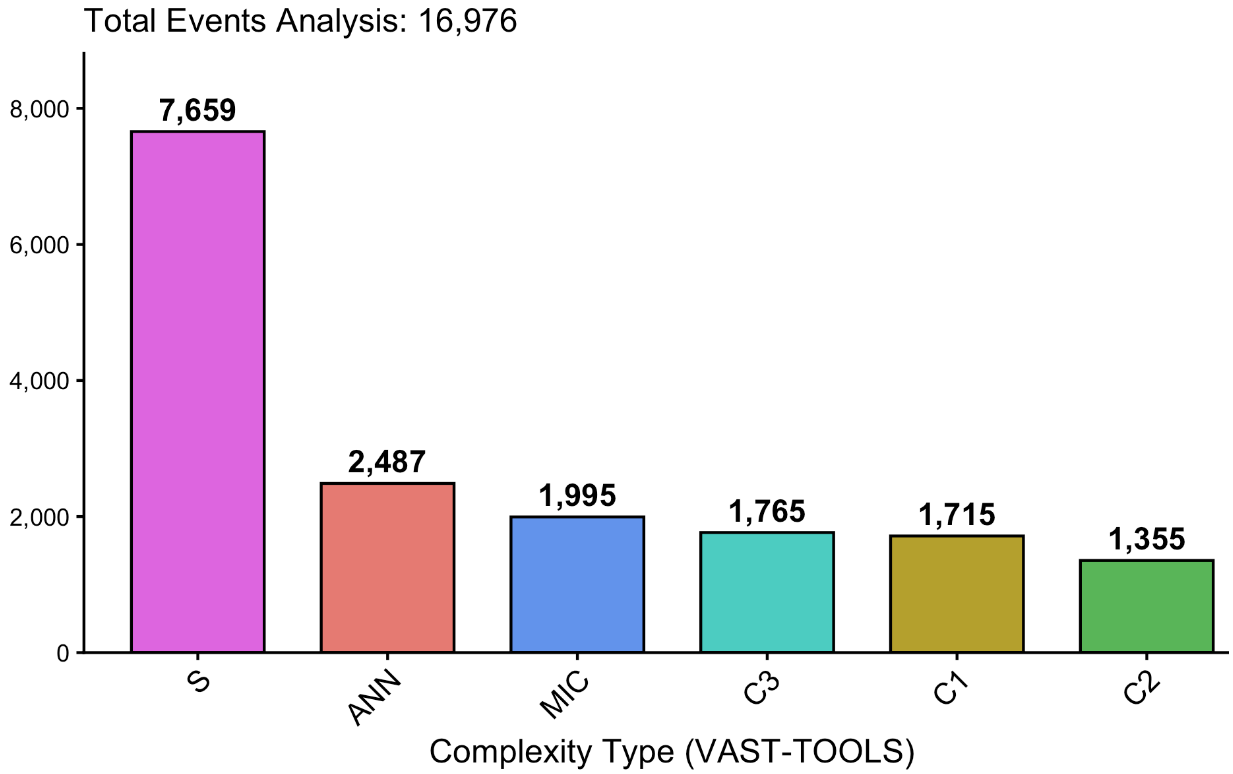


**Total number of events that were considered for the microexon splicing analysis of length upto 51 nts were 16976; where, S: Simple skipping (Standard Alternative Splicing); C1, C2, C3: Increasingly complex events (multiple alternative exons or clusters); IR: Intron Retention; Alt3/Alt5: Alternative 3' or 5' splice sites.All the annotations have been made with respect to the VASTDB**
