## Supplementary Table 1 for "In silico investigation of alternative splicing of microexons in human peripheral tissues"

SUPPLEMENTARY DATA

**Supplementary data table 1:**

**Genes with DSMs found in datasets of the respective diseases with their corresponding ΔPSI values and references indicating their direct or indirect association with the respective disease pathologies:**

| Disease | Gene Name | Microexon event | Coordinate | ΔPSI | Disease Association | | |
|---|---|---|---|---|---|---|---|
| | | | | | Direct | Indirect | Unrelated |
| SCLC | *AP1S2* | HsaEX0004907 | chrX:15828192-15828200 | 68.23885 | ✓ | | |
| | *PACSIN2* | HsaEX0045196 | chr22:42880617-42880622 | 67.17900 | ✓ | | |
| | *DLG1* | HsaEX0019877 | chr3:197075837-197075870 | 80.97549 | | | ✓ |
| | *TMEM87A* | HsaEX1041332 | chr15:42231844-42231887 | -69.30467 | | | ✓ |
| | *CLIP1* | HsaEX0015671 | chr12:122351111-122351143 | -62.01667 | | | ✓ |
| IPF | *ADAM28* | HsaEX1034299 | chr8:24300885-24300929 | 75.00000 | | | ✓ |
| | *TMEM71* | HsaEX1041318 | chr8:132704852-132704901 | 65.30829 | | | ✓ |
| | *FOXP3* | HsaEX0026289 | chrX:49257664-49257699 | 32.6540278 | | | ✓ |
| | *CPNE7* | HsaEX0017013 | chr16:89587556-89587582 | -63.47300 | ✓ | | |
| | *HNF4G* | HsaEX0030264 | chr8:75485785-75485816 | -62.88786 | | | ✓ |
| | *TENM1* | HsaEX0044622 | chrX:124487209-124487229 | -48.291143 | | | ✓ |
| COPD | *ANK2* | HsaEX0004134 | chr4:113348276-113348308 | 26.39152 | | ✓ | |
| | *DMD* | HsaEX1011797 | chrX:32342832-32342850 | 19.88991 | ✓ | | |

| | | | | | | | |
|---|---|---|---|---|---|---|---|
| COPD | *NCAM1* | HsaEX0041923 | chr11:113221296-113221325 | -31.86916 | | ✓ | |
| | *EGFLAM* | HsaEX0021813 | chr5:38445679-38445702 | -20.02333 | ✓ | | |
| COVID | *NCAM1* | HsaEX0041918 | chr11:113236301-113236315 | 66.42750 | ✓ | | |
| | *CHRD* | HsaEX0015287 | chr3:184381080-184381130 | 60.51609 | | ✓ | |
| | *MYH14* | HsaEX0040956 | chr19:50224154-50224177 | 61.72260 | ✓ | | |
| | *CTC1* | HsaEX0017332 | chr19:18744108-18744155 | -73.13286 | | | ✓ |
| | *FSD1L* | HsaEX0026491 | chr9:105513599-105513631 | -51.87019 | | | ✓ |
| HCC | *PRUNE2* | HsaEX0050512 | chr9:76641972-76641980 | 55.18082 | ✓ | | |
| | *FAM13A* | HsaEX0023993 | chr4:89056938-89056988 | 34.04924901 | | ✓ | |
| | *AP3S1* | HsaEX1004151 | chr5:115867805-115867835 | -42.06818 | ✓ | | |
| | *USO1* | HsaEX0069561 | chr4:75795336-75795356 | -42.05077 | ✓ | | |
| | *NDUFAF2* | HsaEX0042301 | chr5:61111762-61111765 | 45.84222 | | ✓ | |
| | *USP21* | HsaEX0069640 | chr1:161160366-161160413 | 34.92279720 | ✓ | | |
| | *TRERF1* | HsaEX0067005 | chr6:42263320-42263365 | -63.33364 | ✓ | | |
| | *MAP3K9* | HsaEX0037611 | chr14:70733757-70733798 | -54.58935 | ✓ | | |
| | *IMMP1L* | HsaEX0031574 | chr11:31469758-31469797 | 81.03000 | | | ✓ |
| | *NACC1* | HsaEX1025044 | chr19:13118419-13118454 | 40.20400 | | | ✓ |

| | | | | | | | |
|---|---|---|---|---|---|---|---|
| DN | *UPF3B* | HsaEX0069453 | chrX:119840646-119840684 | 29.69800000 | | | ✓ |
| | *ZBTB14* | HsaEX0072652 | chr18:5293972-5294001 | -50.14200 | | ✓ | |
| | *MAP3K6* | HsaEX0037585 | chr1:27364661-27364684 | -41.1320000 | | | ✓ |
| LC | *TEAD1* | HsaEX0064361 | chr11:12878889-12878900 | 24.61450549 | | ✓ | |
| | *BAZ2B* | HsaEX0007736 | chr2:159397074-159397100 | 22.04321429 | | | ✓ |
| | *GUCY1A2* | HsaEX1016395 | chr11:107015562-107015599 | -38.37889 | | | ✓ |
| | *ALPK3* | HsaEX0003819 | chr15:84823330-84823368 | -34.35388889 | | | ✓ |
| CH | *DDX11* | HsaEX1011239 | chr12:9288202-9288226 | 32.06275 | | ✓ | |
| | *KSR1* | HsaEX0035062 | chr17:27601360-27601401 | 31.41714286 | | | ✓ |
| | *PRC1* | HsaEX0049880 | chr15:90969079-90969117 | -63.98857 | | | ✓ |
| | *ERBB2* | HsaEX0022931 | chr17:39699537-3969958 | -52.81967 | | ✓ | |
| NAFLD | *PTK2* | HsaEX0050846 | chr8:140669727-140669735 | 41.06492 | ✓ | | |
| | *MYO9B* | HsaEX0041297 | chr19:17210333-17210380 | 25.8217143 | ✓ | | |
| | *DDX11* | HsaEX1011234 | chr12:31091037-31091081 | -67.70917 | | ✓ | |
| | *KCNMA1* | HsaEX1018406 | chr10:77012519-77012557 | -58.98317 | | | ✓ |
| | *XKR8* | HsaEX0071732 | chr1:27964185-27964235 | 60.769375 | ✓ | | |
| RCC | *ABI3BP* | HsaEX0000680 | chr3:100818525-100818569 | 47.89190476 | | ✓ | |

| | | | | | | | |
|---|---|---|---|---|---|---|---|
| RCC | *BAZ2B* | HsaEX0007736 | chr2:159397074-159397100 | -57.01238095 | ✓ | | |
| | *ZBTB14* | HsaEX0072652 | chr18:5293972-5294001 | -44.97798701 | ✓ | | |
| DNE | *COG1* | HsaEX0016177 | chr17:73207702-73207739 | 64.306 | | | ✓ |
| | *CPNE7* | HsaEX0017013 | chr16:89587556-89587582 | 31.690568 | | | ✓ |
| | *SCGB2B2* | HsaEX1036159 | chr19:34632598-34632640 | -42.32166666 | | ✓ | |
| | *ADA* | HsaEX0002246 | chr20:44623034-44623078 | -44.46273148 | ✓ | | |
| CC | *COLQ* | HsaEX0016707 | chr3:15478977-15479003 | 43.75 | | | ✓ |
| | *DMD* | HsaEX1011797 | chrX:32342832-32342850 | 31.89272727 | ✓ | | |
| | *TONSL-AS1* | HsaEX1041607 | chr8:144437699-144437746 | -62.723 | | | ✓ |
| | *CTAGE5* | HsaEX0017658 | chr14:39292680-39292706 | -43.36326797 | ✓ | | |
| UC | *ALG11* | HsaEX0003663 | chr13:52023738-52023779 | 45 | | ✓ | |
| | *ZNF226* | HsaEX7010553 | chr19:44165588-44165613 | 30.8418518 | | ✓ | |
| | *ZIK1* | HsaEX0072812 | chr19:57584952-57584990 | -83.062 | ✓ | | |
| | *CCNE1* | HsaEX0013632 | chr19:29812532-29812578 | -43.32384615 | | ✓ | |
